# Analyzing minimum viable populations in deterministic community models using viability space decomposition

**DOI:** 10.64898/2026.05.19.726018

**Authors:** Eden J. Forbes, Connor McShaffrey

## Abstract

Minimum viable populations (MVPs) are population levels large enough to surmount risk from demographic, environmental, and genetic stochasticity. MVPs are estimated by biologists to guide conservation practices. However, MVPs are generally estimated for a target population without regard for how they interact with intra- and inter-species population dynamics in the broader ecological community. Thus, how and why population dynamics interact with MVPs imposed by conservation biologists remain unclear. When MVPs are imposed on a continuous population model, traditional analyses fail to capture the range of possible outcomes those MVPs create. Here, we describe viability space decomposition (VSD) as a mathematical tool to systematically analyze the potential crossing of MVPs during population dynamics. We demonstrate that different extinction and survival outcomes can be recovered from a model with imposed MVPs using three VSD concepts in junction with a traditional phase portrait: mortality manifolds which separate conditions that lead to different existential outcomes, ordering manifolds which determine the order of extinction events for multiple populations, and collapse manifolds which determine the survival or extinction of one species given the loss of another. We employ these methods with a standard consumer-resource model, and the methods can be scaled to systems with more species. VSD is a useful tool for conservation biologists and community ecologists concerned with boundary crossing problems in any dynamical system.

## INTRODUCTION

Small populations are more likely to go extinct due to demographic, environmental, and genetic stochasticity (Nunney and Campbell 1993, Melbourne and Hastings 2008). Minimal viable populations (MVPs) are population thresholds at which these stochastic forces no longer pose a threat of extinction (Shaffer 1981, Gabriel and Bürger 1992). MVPs are regularly invoked in contemporary conservation biology and ecology for both theory and practice (e.g., Courchamp et al. 2018, Williams et al. 2021, Horswill et al. 2022, Pérez-Pereira et al. 2022, Mills et al. 2025). That said, MVPs are generally defined or modeled in reference to a focal species rather than in the context of a whole community. While MVPs are a well-known concept, mathematical tools are limited in how they can comprehensively inform management decisions in a given ecological community. No current body of work, to our knowledge, describes how population dynamics interact with MVPs or how interventions impact those dynamics given nonlinear interactions among populations. Even after the arduous experimental and mathematical work has approximated an MVP (Coulson et al. 2001, Reed et al. 2002, Traill et al. 2007, Flather et al. 2011), that MVP’s role relative to community dynamics remains unclear.

How and why do MVPs change the predicted outcomes of community population dynamics? Conversely, how and why do community dynamics change the predicted outcomes of imposing MVPs? Community models commonly only represent species extinction at zero population, allowing for exceedingly small populations that may be unrealistic in natural systems. To account for the precarity of small populations, ecological models often add biological assumptions that generate density-dependent reproduction. Most frequently, Allee effects, in which a species experiences an increase in growth rate with an increase in its own density (Allee 1938), are employed to represent the extinction risk for low-density populations (Lande 1987, Dennis 1989). Strong Allee effects in particular yield negative growth rates for small populations. As such, Allee effects have been deemed an important factor to consider for conservation (Stephens and Sutherland 1999, Courchamp et al. 2008) and may be assumed as the organizing structures underlying MVPs in community models. That said, Allee effects do not encompass the stochastic processes underlying empirical MVP estimates (Stephens et al. 1999). An MVP should be defined as a population that cannot be perturbed to extinction by forces within an expected range of intensity, whether that population be above the Allee threshold (and expected to survive) or below it (and expected to collapse). Therefore, while Allee effects may contribute to MVPs and shape their influence on an ecological community, they are not a mathematical stand-in for MVPs.

We posit that MVP research in the context of ecological communities needs new mathematical tools to understand how population dynamics will interact with an estimated MVP. In this manuscript, we demonstrate how viability theory is an effective tool for the treatment of MVPs in ecological modeling. Viability theory describes systems that can only persist if they remain within biophysical constraints called *viability constraints* (Ashby 1960). Recently, numerous tools for viability analysis have been developed from dynamical (Barandiaran and Egbert 2014), information-theoretic (Egbert and Pérez-Mercader 2018, Kolchinsky and Wolpert 2018), and control-theoretic (Aubin 1991, Himeoka et al. 2024) perspectives. Notably, though, the models examined in each of these frameworks usually only involve the persistence of a singular organism (Barandiaran and Egbert 2014, Kolezhitskiy et al. 2022, McShaffrey and Beer 2022) or system (Cury et al. 2005, Aubin and Saint-Pierre 2007) rather than two or more interacting entities. A method that identifies global structures in state space at such a scale is needed. For this reason, we explored MVPs from the perspective of *viability space decomposition* (VSD), a framework developed for single- and multi-organism ODE models that identifies the structures which globally decompose state space into regions of distinct survival outcomes for multiple systems (here, populations; McShaffrey and Beer 2023, 2024, 2026). Just like the physiological constraints of individual organisms, we show that MVPs can be represented as the viability boundaries of whole populations whose violation puts that population at risk of extinction. VSD can thus determine the sequence of potential extirpation events as a function of the current state of a modeled community.

To demonstrate the utility of VSD in ecological models, we examined a simple consumer-resource model with MVPs imposed on both species. The consumer clears the resource according to a type III functional response while the resource grows cubically, yielding a strong Allee effect with and without the consumer (Kot 2001). Consequently, we explored the distinction between and potential interaction of MVPs and Allee effects directly. As parameterized, the C-R system without MVPs demonstrated two stable regimes: mutual survival and mutual extinction. We then simulated thousands of initial conditions to determine how model outcomes changed with MVPs imposed, assuming violation of the MVP yields local extinction (a high-risk environment). Those simulations revealed four possible outcomes for the C-R system, none of which were fully explained by traditional dynamical analyses for ecological models. We then demonstrated how VSD recovers the structure separating those different outcomes. VSD can thus help explore how community dynamics interact with measured MVPs, as well as the consequences of management responses maintaining populations relative to those MVPs.

### MODEL STRUCTURE, BEHAVIOR, AND MINIMAL VIABLE POPULATIONS

Our focal model depicts a two-species consumer-resource system (Table 1; Fig. 1A):

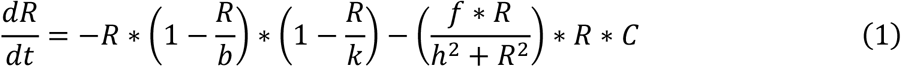

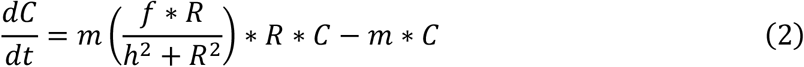

**Table 1:** Parameter definitions and viability space decomposition notation.

| *Term* | *Definition* | *Value/Range* |
| --- | --- | --- |
| **Parameters** |  |  |
| $b$ | Growth rate inflection, resource | 1 |
| $f$ | Feeding rate, consumers | 1.5 |
| $h$ | Half-saturation constant, feeding rate | 1 |
| $k$ | Carrying capacity, resource | 1.6 |
| $m$ | Metabolic rate, consumers | 0.8 |
| $R_{MIN}$ | Minimal viable population density, resource | 0.5 |
| $C_{MIN}$ | Minimal viable population density, consumer | 0.05 |
| **Viability Notation** |  |  |
| $A$ | Asymptotically viable region | - |
| $G_i$ | Guard condition for violating the MVP for species $i$ | - |
| $\mathbf{m}$ | Mortality points | - |
| $M$ | Mortality manifold | - |
| $n_i$ | Orthogonal vector to the MVP for species $i$ | - |
| $\mathbf{o}$ | Ordering points | - |
| $O$ | Ordering manifold | - |
| $T$ | Transiently viable region | - |
| $U$ | State space of all species | - |
| $V_i$ | Viable region of species $i$ | - |
| $x$ | A state, or a consumer-resource population, in $U$ | - |
| $X$ | Intersection of all $V_i$ | - |
| $\Lambda_i$ | Points tangent to the MVP of species $i$ | - |
| $\phi$ | System flow | - |
| $\mathcal{U}_i$ | Collapse function given the satisfaction of $G_i$ | - |

**Figure 1.**
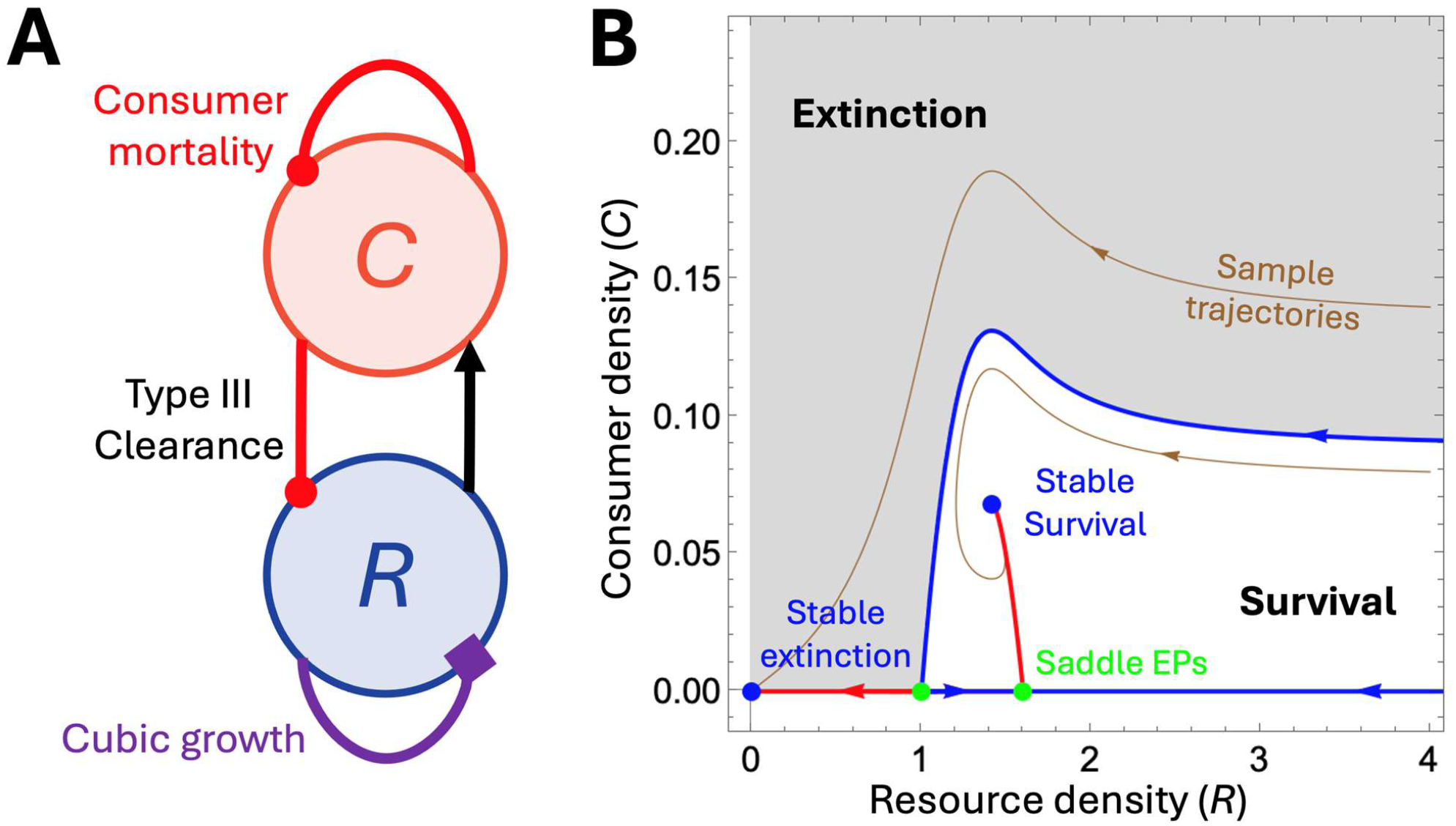
Cubic growth of a resource in a consumer-resource system yields bistability. **(A)** The consumer-resource model is governed by cubic growth of the resource (which can yield either self-inhibition or self-excitation; purple diamond), type III clearance of the resource by the consumer (excitatory for the consumer [black arrow] and inhibitory for the resource [red punch]), and consumer mortality (inhibitory, red punch). **(B)** The consumer-resource system demonstrates an Allee effect with two possible outcomes. At high initial resource density and low initial consumer density, both populations converge on the stable survival equilibrium. At low initial resource densities and low initial consumer densities, the system converges on the simultaneous extinction of both species. Blue points indicate stable equilibria, while green points indicate partially unstable saddle equilibria. Stable and unstable saddle manifolds are denoted in blue and red, respectively. Sample trajectories of system behavior are demonstrated in brown.

The resource (*R*; eq. 1) grows cubically up to carrying capacity *k*, with positive growth rates at densities above an inflection point determined by density *b* (where *b < k*) and negative growth rate below that density. Resources thus can establish stable populations with a large enough initial population but fail to do so if the population becomes too small (falls below *b*). So, the resource *R* demonstrates an Allee effect independent of the consumer, with the possibility of either stable persistence at carrying capacity *k* or stable extinction at *R = 0*. Resources are cleared by consumers according to a type III functional response (Holling 1959) governed by maximum consumer feeding rate *f* and a half-saturation constant *h* denoting when feeding rate reaches half that maximum. The consumer population (*C*, eq. 2) in turn grows according to that clearance function. Both conversion of cleared resources and consumer metabolic loss (thus, mortality) are scaled by consumer metabolic rate *m* (like in McCann et al. 1998).

Like the resource alone, the C-R model demonstrates a strong Allee effect, or bistability, between a stable C-R state and an extinction state (Fig. 1B). Whether or not an initial set of population densities falls into one or the other state can be determined by examining the model’s basins of attraction. Initial conditions that yield persistent populations (white area, Fig. 1B) and those that yield extinction (gray area, Fig. 1B) are separated by a stable saddle manifold (blue line, Fig. 1B) which extends from the unstable saddle equilibrium point determined by *b*. The previously stable point where the resource alone is at carrying capacity *k* now is a saddle with an unstable manifold (red line, Fig. 1B) extending to the new stable C-R equilibrium. If we accepted the Allee effect as a stand-in for the resource’s MVP (*R*_*MIN*_ *= b*), the stable saddle manifold that separates the two basins would determine whether a given initial set of populations would lead to survival or extinction. However, the base C-R model allows for vanishingly small consumer densities which may be at risk from demographic, ecological, and genetic stochasticity, or single-time perturbations (Shaffer 1981). As such, we imposed an MVP on both the resource (*R*_*MIN*_ = 0.5) and the consumer (*C*_*MIN*_ = 0.05). Assuming the worst possible case scenario, species were considered locally extinct if their population violated an MVP, thus dynamics collapse to a lower dimensional community model with the remaining species. This assumption is relevant if (1) there is high environmental risk, (2) stochasticity dominates population dynamics at small population sizes, or (3) any amount of risk beyond the MVP is unacceptable to managers

We simulated population dynamics for large ranges of initial population densities above both MVPs (*R* [0.5: 5]; *C* [0.05: 0.1]) and examined if and when each species fell below its MVP and the order in which those violations occurred. Imposing MVPs on the C-R model introduces new regimes beyond mutual survival and simultaneous extinction (Fig. 2). Now, either the resource or consumer can be the first to go locally extinct (dark and medium gray regions, respectively), the resource can survive in the absence of the consumer (light gray region), or both species can persist (white region). Critically, traditional dynamical analysis no longer explains these outcomes, as the stable manifold that previously separated survival and extinction no longer correlates with the simulated outcomes. As such, relying on the standard dynamical analysis of an Allee effect would not predict the behavior of the system with imposed MVPs. Instead, additional techniques are required to derive the consequences of MVPs analytically rather than relying on computationally expensive simulations.

**Figure 2.**
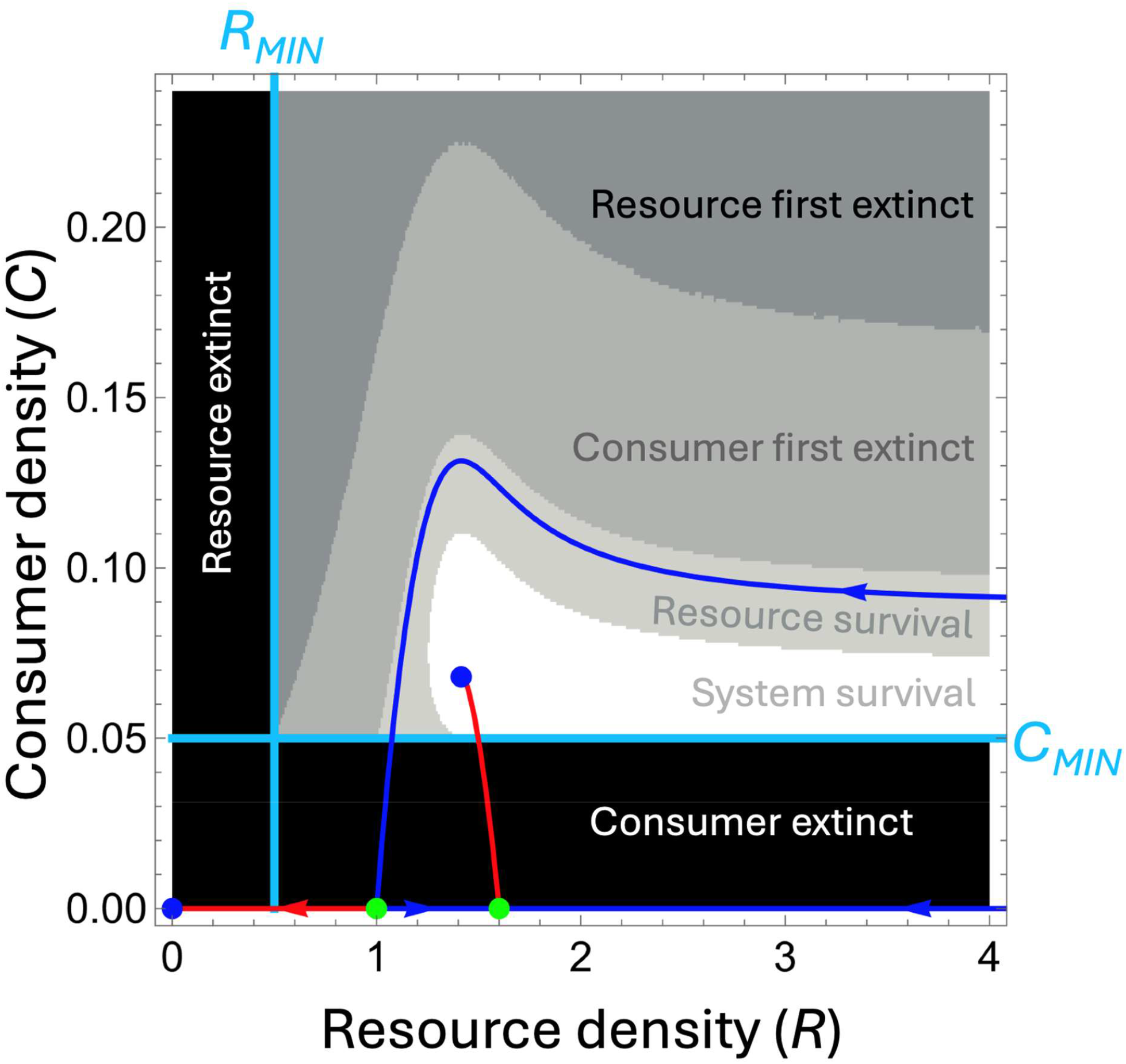
Minimal viable population (MVP) constraints introduce new unexplained system behaviors. MVP constraints (light blue) define regions of extinction risk (here assume to be instantaneous extinction) with positive densities. Simulations of the system with these constraints yield four possible outcomes: either the resource goes extinct followed by the consumer (darkest gray), the consumer goes extinct followed by the resource (medium gray), the resource survives but the consumer goes extinct (light gray), or both species survive (white). Equilibria follow the conventions in Fig. 1. Notably, boundaries between the order and occurrence of extinctions no longer coincide with any of the traditional dynamical manifolds.

### VIABILITY SPACE DECOMPOSITION

Characterizing the role of MVPs in our C-R model requires an extension of dynamical systems theory that describes how model dynamics (the two-species model) interact with imposed boundaries (MVPs). To this end, we used ‘viability space decomposition’ (VSD), a theory that describes new global structures in dynamical systems with boundary crossing elements (McShaffrey and Beer 2026). In our case, these boundaries are *R*_*MIN*_ and *C*_*MIN*_, the MVPs for resource and consumers, respectively. We assumed that all population trajectories either converged to a viable attractor or transiently violated an MVP. We assumed that if a population violates its MVP, it is extinguished instantly rather than applying some probability of extinction (although for a treatment of VSD on probabilistic models, see McShaffrey and Beer 2025). As such, we assumed that trajectories of the two-species system do not extend beyond the MVP limits *R*_*MIN*_ and *C*_*MIN*_. We present three classes of global manifolds described by VSD (‘mortality’, ‘ordering’, and ‘collapse’) that each can be derived to divide up the outcomes demonstrated in Fig. 2.

#### Viability space decomposition definitions

Defining viability constraints (here, MVPs) converts a fully continuous dynamical system (F(*x*), in our case eq. 1 and 2) into a hybrid system that combines continuous dynamics with discrete transitions between modes (population existence and extinction). Given a state space *U* that comprises all the populations currently intact, the viability region *V* of each population is defined relative to its respective MVP:

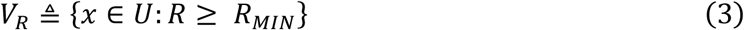

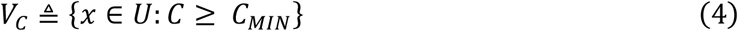

Extinction occurs at the single moment when an MVP is violated. Thus, in this case a population’s dynamics are undefined outside of its viability region such that the flow *ϕ* does not cross the boundary of *V*:

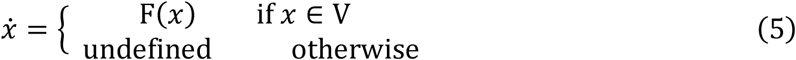

Thus, when both populations are present, the dynamics can only unfold without extinction events in the intersection of their viability regions:

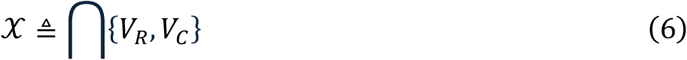

Notably, because dynamics are only defined relative to the intersection of viability regions, of the limit sets (Fig. 1B) only the stable equilibrium point exists within 𝒳. The flow that would have been generated by F(*x*) beyond this region is, in a sense, “virtual,” in that it depicts dynamics that would have been possible under different viability constraints. Relatedly, the virtual saddle node in the complement of 𝒳 is not a meaningful organizing structure for this model’s combination of change equations and constraints. The trajectory segment in 𝒳 that would have corresponded to that saddle node’s stable manifold becomes just another trajectory *as opposed to* an organizing boundary. Understanding what new global structures take the stable manifold’s place in organizing our flow requires that we think critically about how the continuous dynamics and viability boundaries interact.

Explicitly representing extinction in a hybrid dynamical system requires two additional features based on viability regions. The first is a guard condition *G*_*i*_ that specifies a population is only extinguished if it meets its MVP, the boundary of the viability region, with an outward velocity (purple trajectories, Fig. 3A):

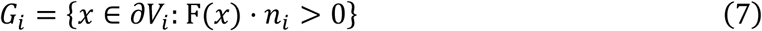

**Figure 3.**
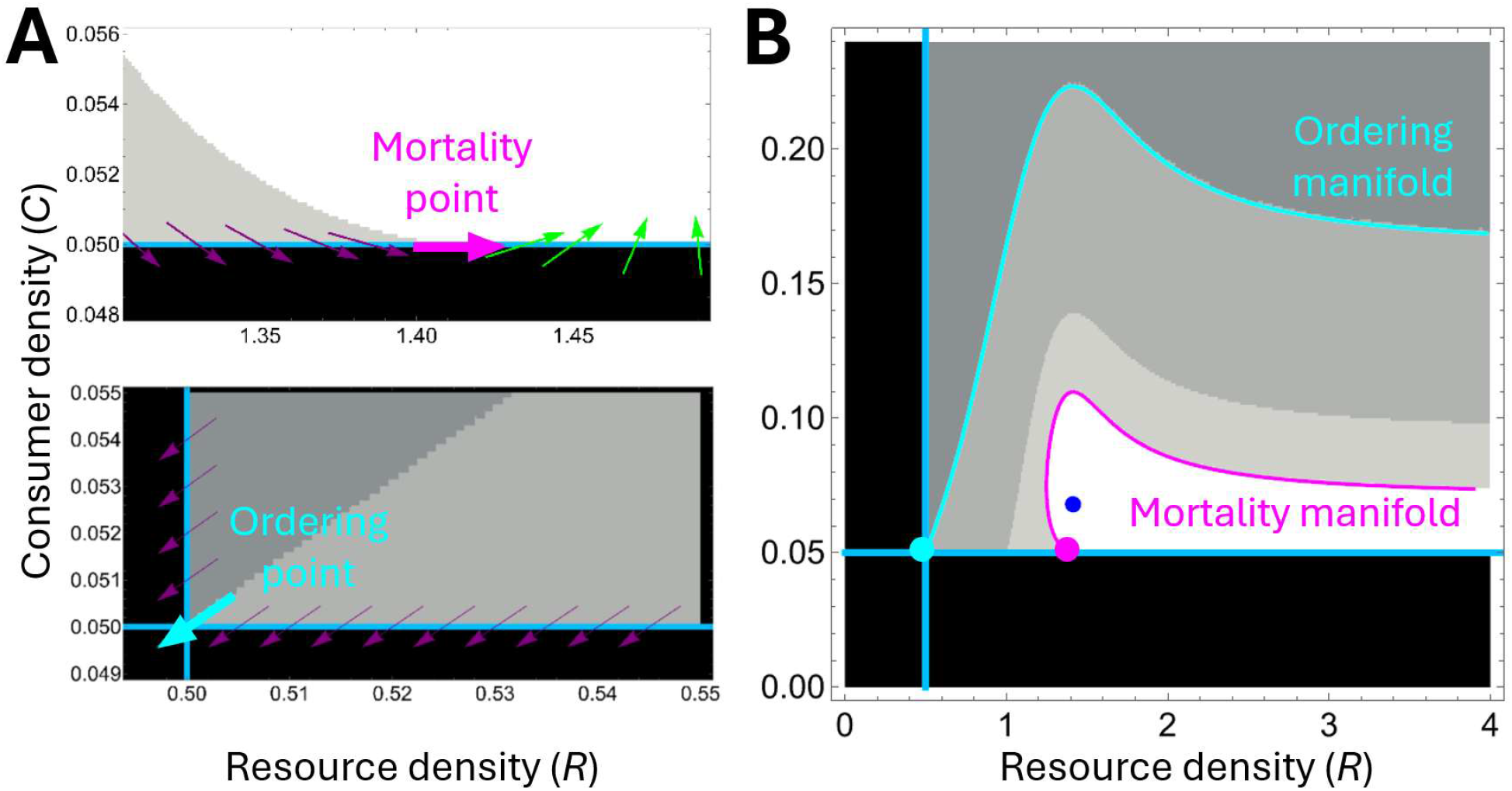
**(A)** Mortality manifolds delineate whether both consumer and resource populations are asymptotically viable/will survive. Mortality manifolds are determined by simulating the backwards time trajectory of points tangent to a given MVP (‘mortality points’; magenta arrow) which separate points along the MVP that leave the viable region (purple arrows) from those which remain viable (green arrows). Ordering manifolds delineate which population will collapse first when both species are transiently viable. Like mortality manifolds, ordering manifolds are determined by simulating the backwards time trajectory of points where MVPs (blue lines) intersect (‘ordering point’; cyan arrow). **(B)** Here, a mortality manifold exactly separates cases in which both consumers and resources are asymptotically viable and cases in which only resources are asymptotically viable. An ordering manifold exactly separates cases in which resources are first extinguished and consumers are first extinguished, given both are transiently viable.

where *n*_*i*_ is the vector orthogonal to the viability boundary of population *i* that points into the complement of the viability region:

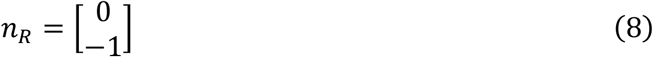

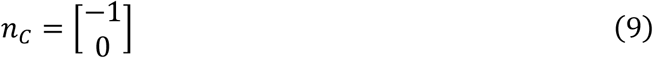

Second, we need a reset function that specifies how the system should change after the guard conditions have been met. In our case, this will be a collapse function (⇓_*i*_) that describes the extinction of a population *i* and thus the removal of its state dimension as a discrete event. For example, given our system is comprised of only two species, the extinction of the consumer suggests:

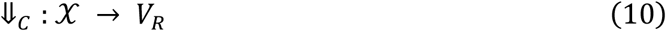

Notably, if multiple guard conditions are satisfied at once, the corresponding collapse functions can be composed to remove the state dimensions of all the populations being lost in the extinction event.

#### Mortality and ordering manifolds

With the hybrid system fully defined (eq. 1 – 10) by the continuous system F(*x*), MVPs, resulting viability regions, guard conditions *G*_*i*_, and collapse functions ⇓_*i*_, we can separate initial conditions in any intersection of viability regions or single viability region into two general classes, those that are *asymptotically* viable (*A*) and those that are *transiently* viable (*T*), where we only consider initial conditions at the MVPs or above:

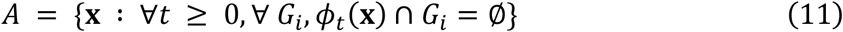

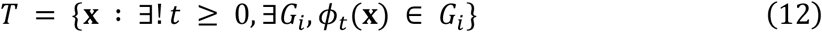

Asymptotically viable trajectories are those that never violate an MVP. Generally, *A* requires at least one viable attractor to which asymptotically viable trajectories will converge. Conversely, transiently viable trajectories are those that eventually violate an MVP. Thus, they indicate cases in which one or more species exist but are on their way to extinction. Importantly, these sets only characterize the survival outcomes for the current set of populations; additional steps are required to characterize what happens after the first extinction event, and we will show these later.

As discussed, traditional dynamical systems analysis does not cleanly separate *A* and *T* for either species in our model (Fig. 2). Instead, we need a more focused treatment of how F(x) interacts with a given MVP. Taking *C*_*MIN*_ as an example, we can see that the two most common change vectors at an MVP are either recovery vectors that move towards the interior of 𝒳 (green arrows, Fig. 3A) and fatality vectors characterized by *G*_*i*_ (eq. 7) that have outward velocity (purple arrows, Fig. 3A). However, smoothly transitioning between recovery and fatality vectors required we pass through a point where the change vector is tangential to the MVP. The set of tangent points Λ_*C*_ thus separates recovery and fatality points on the MVP:

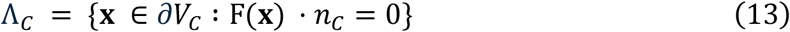

To carve out subsets of the viability regions with different fates, we needed to further separate the tangent points along an MVP into two sets: those that eventually violate that same MVP, and those that converge to a different MVP or an attractor. The latter set of tangencies are referred to as *mortality points* (**m**). We find one such tangency which converges to our stable equilibrium:

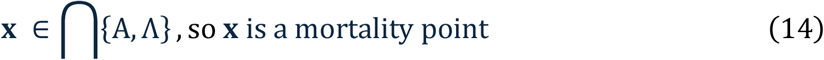

The backward time trajectories of these mortality points provide mortality manifolds (*M*) which separate initial conditions based on which of the two aforementioned fates to which they commit:

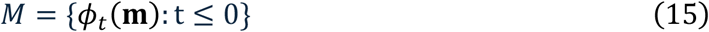

In our C-R model, we identified no mortality points on *R*_*MIN*_ but did find one mortality point on *C*_*MIN*_. The mortality manifold generated from this point (magenta curve, Fig. 3B) exactly separated conditions in which both species survived and those where only the resource survived in our simulations.

If both of our species are transiently viable, that is, on the path to extinction, we can use a similar technique to determine the order in which those species will be put at risk. To do so, we must identify *ordering points* (**o**) where both species’ guard conditions *G*_*i*_ (eq. 7) are met simultaneously:

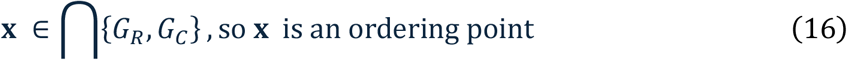

Like mortality points, ordering points can be integrated backward in time to create *ordering manifolds*, which separate two transiently viable regions by which MVP is violated first:

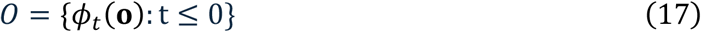

In our two-species model, we identified one ordering point where *R*_*MIN*_ and *C*_*MIN*_ intersected. The ordering manifold generated from this point (cyan curve, Fig. 3B) exactly separated conditions in which the resource *R* was first put at extinction risk and those in which the consumer *C* was first put at extinction risk.

#### Collapse manifolds

So far, mortality and ordering manifolds have recovered two of the three separatrices evident in our simulated outcomes. Lastly, we need an account of what distinguishes whether the resource will end up asymptotically viable or transiently viable once the consumer has gone extinct (‘consumer first extinct’ vs. ‘resource survival’, Fig. 2). Ultimately, the outcome for the resource depends on where the collapse function ⇓_C_ (eq. 10) lands the population in its isolated viability region *V*_*R*_. Similar to how we found mortality and ordering manifolds by integrating relevant points backwards in time, the collapse function can be inverted. Because a species’ collapse function is only triggered when that species guard condition *G*_*i*_ is satisfied (eq. 7), the inverse of the collapse function maps from the lower dimensional viability region (in this case *V*_*R*_) to points on the boundary triggering the collapse function (in this case *C*_*MIN*_, thus collapse function ⇓_C_):

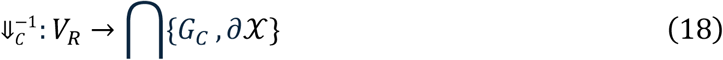

Thus, if we have an organizing manifold in a lower dimension, the inverse of the collapse function can map which terminal points along the violated boundary collapse exactly onto that manifold. In the case of the resource, the organizing manifold is just the unstable equilibrium created by the cubic growth function (in eq. 1) which separates resource extinction via violation of *R*_*MIN*_ and persistence at the upper stable equilibrium point (Fig. 4A). As before, we calculated the backwards time trajectory of the point created by joining that unstable equilibrium on the x-axis with *C*_*MIN*_ on the y-axis. The resulting collapse manifold (red dashed curve, Fig. 4B) exactly separated our final two regions, determining which conditions in C-R space would lead to survival or extinction of the resource given the consumer species was transiently viable (determined by the mortality manifold). Thus, by measuring mortality, ordering, and collapse manifolds, we have fully detailed the various survival outcomes of the C-R system with MVPs (Fig. 5).

**Figure 4.**
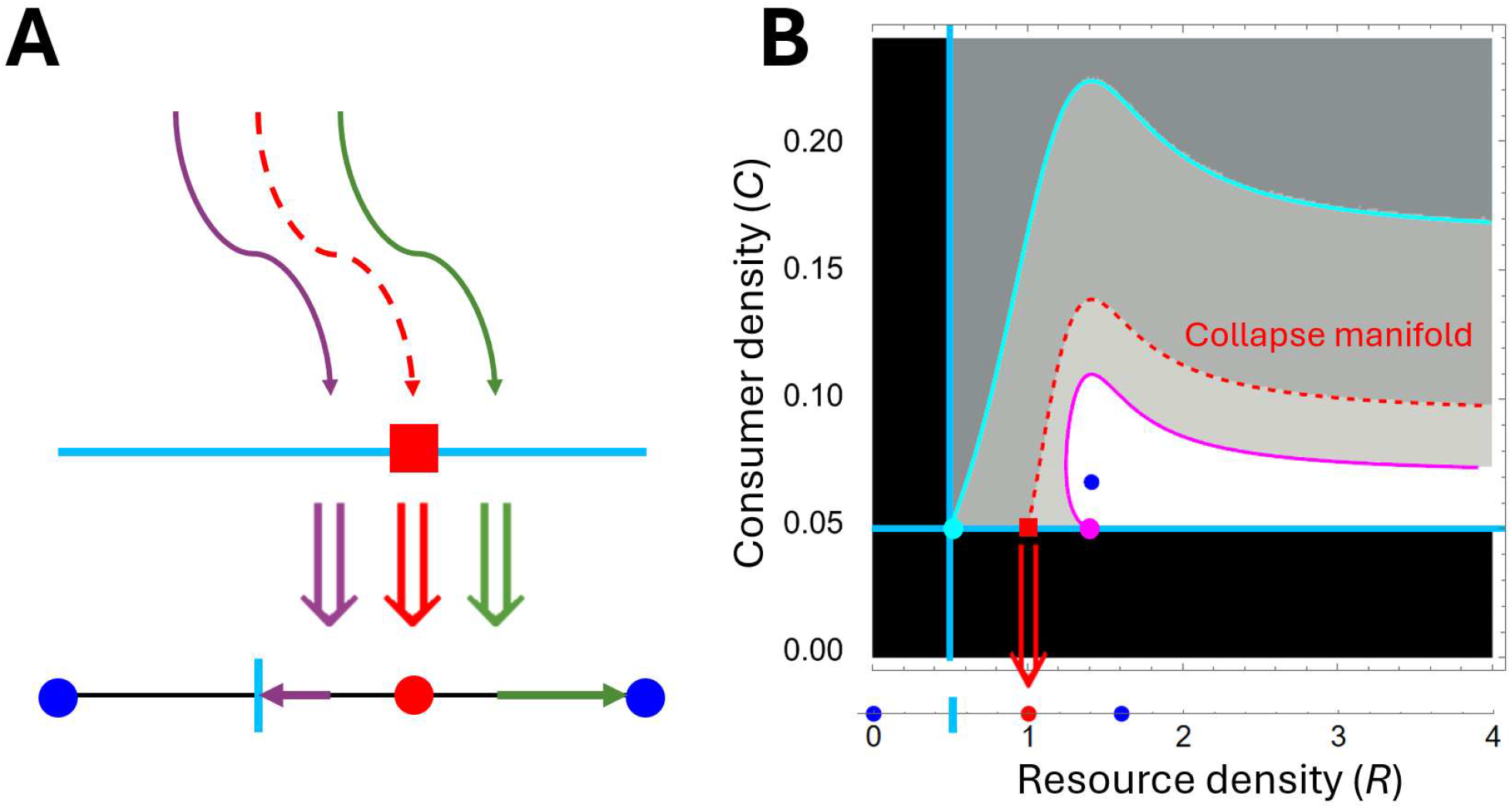
Collapse manifolds determine the viability of populations when another local population goes extinct. **(A)** When an MVP is violated, the remaining species in the community are defined by a system one dimension lower than before. Initial conditions in the new system are determined by population densities when the MVP was violated. Where an MVP is violated can determine outcomes in the lower dimensional system. **(B)** The collapse manifold (red dashed line) separates cases in which the resource is asymptotically viable versus transiently viable.

**Figure 5.**
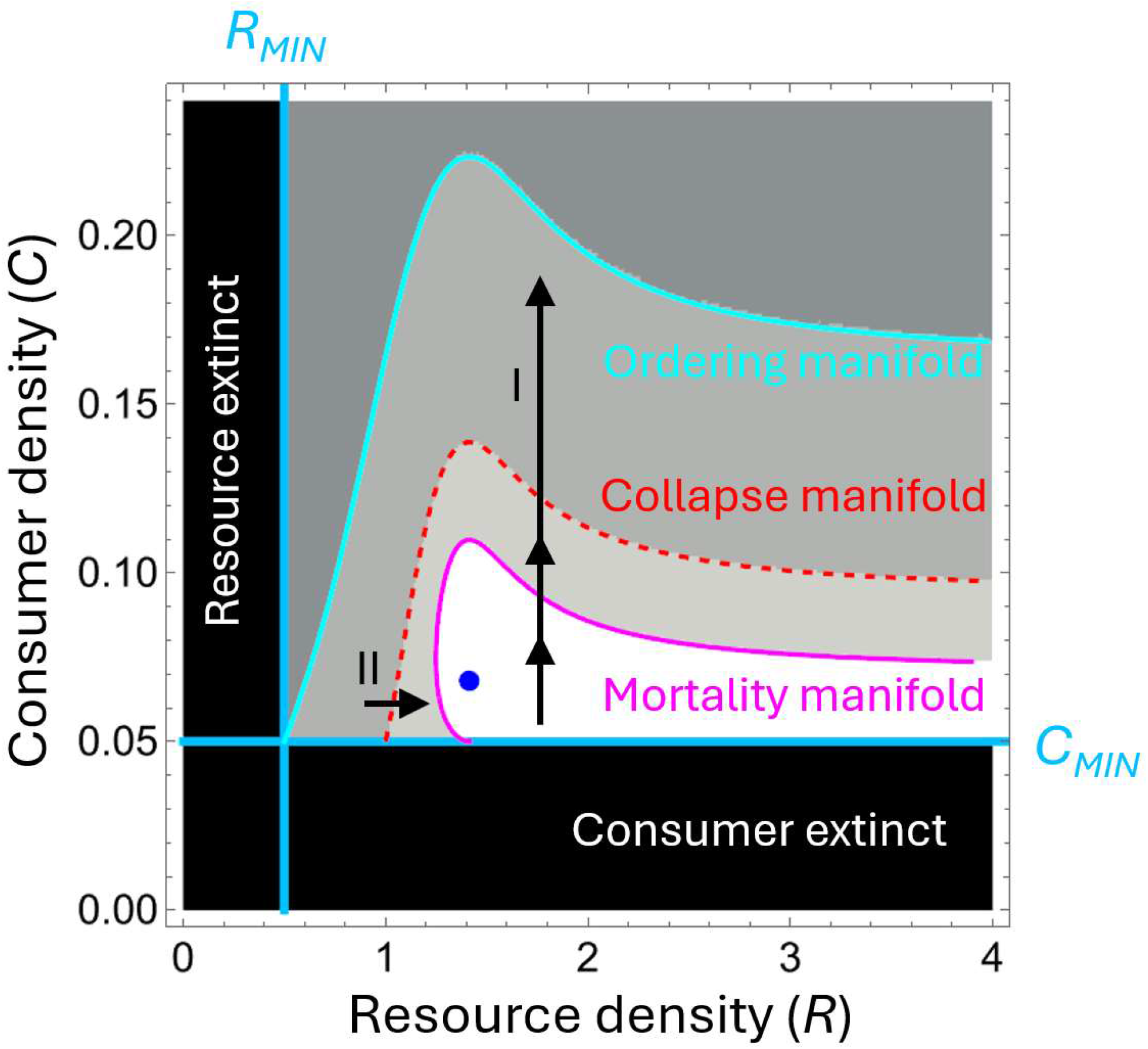
In conjunction, viability space decomposition (VSD) identified mortality (magenta), ordering (cyan), and collapse (red dashed) manifolds that fully explain the various survival outcomes in the consumer-resource model with imposed MVPs. Perturbations to the system (e.g., via management strategies) are now interpreted differently than before employing VSD. For example (I), consumer overstocking can not only push a consumer to extinction (crossing a mortality manifold; magenta line) but can also push the resource to extinction (crossing a collapse manifold; cyan line) and cause the resource to collapse first. Conversely, (II) even when management perturbations cannot rescue a community, they may still be able to rescue certain species (crossing collapse manifolds; red dashed line).

## DISCUSSION

We have demonstrated viability space decomposition (VSD) as a method for applying minimum viable population (MVP) thresholds to standard community models in ecology. While traditional dynamical analyses fail to account for the survival outcomes of population models with thresholding limits such as MVPs, VSD recovers those outcomes exactly. Mortality manifolds separate regions where initial populations will all remain viable and those where one or more species will violate an MVP. Ordering manifolds describe which species will first violate an MVP in circumstances where multiple species are transiently viable. Collapse manifolds describe the downstream consequences of a local extinction for the remaining species in a community, namely whether those remaining species will stabilize or be exposed to further risk. Furthermore, VSD is relatively simple to employ and requires negligible computational time, especially compared to the computationally expensive brute-force simulation approach we used to initially generate the range of C-R outcomes with imposed MVPs. Scaling still has its limits, as approximating mortality, ordering, and collapse manifolds is more challenging with more species. Still, even knowing the existence of these manifolds can support more efficient numerical exploration of more diverse community models.

VSD also allows for more nuanced connections between community models and management decisions. Several examples fall out from this C-R model. First, overstocking of the consumer species in the C-R system without MVPs could lead to extinction risk for both species when the resource was subject to an Allee effect. VSD demonstrates how consequences of overstocking can reverberate throughout a community (e.g. Ims et al. 2007), predicting both whether the resource may be put at risk and in which order consumers and resources may collapse (trajectory I, Fig. 5). Second, assume that a certain management strategy may only achieve a small change in a population (trajectory II, Fig. 5). VSD demonstrates whether those small changes may still prevent secondary extinction risk or alter the sequence of extinctions even if the consumer species cannot be conserved (Eklöf and Ebenman 2006, Dunne and Williams 2009, Brodie et al. 2014). Third, comparing VSDs for different MVPs in a model may suggest different management strategies, given MVPs may change in definition (different levels of risk) or in value (via more systemic changes to an ecosystem or longer-term changes in life history, e.g. Wang et al. 2019). In all cases, synthesizing MVPs with population dynamics demands specific claims about life history and environmental context that change the role of a purported MVP in management. Synthesis of MVPs with population dynamics is an important step given that MVPs and efforts in response to those MVPs are often erroneous without appropriate context (Flather et al. 2011).

We assumed that MVPs had been identified for the modeled populations and that stochastic processes were not necessary to include. Nonetheless, even for models that do require noise, deterministic viability analyses such as VSD can still provide important qualitative insights. For example, in cases of additive noise, mortality manifolds can get blurred into fuzzy bands where the norm of the average first-passage time gradient is especially steep, and these bands separate regions where the gradient is relatively shallow (McShaffrey and Beer 2025). Exploring the consequences for communities of MVPs of varying severity (or the steepness of the stochastic shift) is an important future direction for VSD in ecology especially as pertains to demographic (Gabriel and Bürger 1992) and genetic stochasticity (Frankham et al. 2014). Viability boundaries themselves can be defined in a fuzzy and probabilistic manner and while leaving the dynamics deterministic, although this would not fundamentally change the VSD process. For example, if we wanted to identify where there was a 20% chance of extinction, this would be manifold in the space that looks much like our deterministic viability boundaries. We could then solve for the regions of trajectories that would cross the 20% threshold. Therefore, a continuum of VSD manifolds corresponding to different probabilities of extinction would be likely.

While MVPs are one obvious application for VSD in ecological modeling, the same methods apply to any other thresholds imposed onto ecological models. There are several immediate extensions we see for VSD in ecological modeling. Ongoing work is characterizing how VSD objects change as parameters are varied, allowing the study of *viability bifurcations* (McShaffrey and Beer 2023). Such analyses would facilitate VSD analysis across ranges of a given parameter, facilitating sensitivity analysis of VSD objects. Additionally, combining VSD with feedback loop analysis (e.g. Puccia and Levins 1985, Novak et al. 2016, Forbes and Hall 2026, Forbes and Stockwell 2026) could help determine what relationships cause boundary violations at different locations, given that different trophic relationships may drive a given population’s decline at different community states. Other ongoing work is focused on effectively scaling VSD to higher dimensions. Increased dimensionality has import on many VSD objects (e.g., simultaneous extinction of species). Future work should focus on describing paths of invasion and collapse in ecological communities with MVPs which are important in determining the pace of gain and loss in ecosystem function (Larsen et al. 2005, Srinivasan et al. 2007, Rumeu et al. 2017). Finally, VSD applies to other ecological interests as well (e.g. bioenergetics models such as dynamic energy budgets; Kooijman 2009), as it generally describes the consequences of imposing boundaries or thresholds on flows generated by ordinary differential equations.

We have shown that VSD is an effective tool for demonstrating the consequences of researcher-determined (either by measurements of stochasticity or management choice) boundaries on ecological communities. Our results offer new insight into a canonical ecological model and can do the same for theoretical and empirically motivated community models alike.

## ACKNOWLEDGEMENTS

The authors would like to thank Randall D. Beer, members of the Computational Neuroethology Lab at Indiana University, and members of the Rubenstein Ecological Laboratory at University of Vermont for their feedback on earlier versions of this work. C. M. is supported by the National Science Foundation Graduate Research Fellowship Program under Grant No. 2240777. Any opinions, findings, and conclusions or recommendations expressed in this material are those of the author(s) and do not necessarily reflect the views of the National Science Foundation.

## AUTHOR CONTRIBUTIONS

**E. J. F**. Conceptualization (equal), formal analysis (lead), methodology (supporting), software (equal), writing – original draft (equal), writing – review & editing (equal), visualization (lead). **C. M**. Conceptualization (equal), formal analysis (supporting), methodology (lead), software (equal), writing – original draft (equal), writing – review & editing (equal), visualization (supporting)

